# Enabling Precision Medicine via standard communication of HTS provenance, analysis, and results

**DOI:** 10.1101/191783

**Authors:** Gil Alterovitz, Dennis Dean, Carole Goble, Michael R. Crusoe, Stian Soiland-Reyes, Amanda Bell, Anais Hayes, Anita Suresh, Anjan Purkayastha, Charles H. King, Dan Taylor, Elaine Johanson, Elaine E. Thompson, Eric Donaldson, Hiroki Morizono, Hsinyi Tsang, Jeet K. Vora, Jeremy Goecks, Jianchao Yao, Jonas S. Almeida, Jonathon Keeney, KanakaDurga Addepalli, Konstantinos Krampis, Krista M. Smith, Lydia Guo, Mark Walderhaug, Marco Schito, Matthew Ezewudo, Nuria Guimera, Paul Walsh, Robel Kahsay, Srikanth Gottipati, Timothy C Rodwell, Toby Bloom, Yuching Lai, Vahan Simonyan, Raja Mazumder

**Affiliations:** Harvard/MIT Division of Health Sciences and Technology, Harvard Medical School, Boston, MA 02115, USA; Computational Health Informatics Program, Boston Children’s Hospital, Boston, MA 02115, USA; Electrical Engineering and Computer Science, Massachusetts Institute of Technology, Boston, MA 02139, USA; Seven Bridges, Cambridge MA, 02142, USA; School of Computer Science, The University of Manchester, Manchester, M13 9PL, UK; Common Workflow Language Project, Vilnius, Lithuania; The Department of Biochemistry & Molecular Medicine, The George Washington University Medical Center, Washington, DC 20037, USA; Foundation for Innovative New Diagnostics (FIND), Chemin des Mines 9, 1202 Geneva, Switzerland; OpenBox Bio, LLC, USA; The McCormick Genomic and Proteomic Center, The George Washington University, Washington, DC 20037, USA; Internet 2, 1150 18^th^ St. NW, Washington, DC 20036, USA; US Food and Drug Administration, Silver Spring MD 20993, United States of America; Center for Genetic Medicine, Children’s National Medical Center, Washington, DC 20010, USA; The Department of Genomics and Precision Medicine, The George Washington University School of Medicine and Health Sciences, Washington, DC 20037, USA; Center for Biomedical Informatics and Information Technology, National Cancer Institute, National Institutes of Health, Gaithersburg, MD, USA; Attain, LLC, McClean, VA, USA; Computational Biology Program, Oregon Health & Science University, Portland OR, 97239, USA; MRL IT, Merck & Co., Inc., Boston, MA, USA; Stony Brook University, School of Medicine and College of Engineering and Applied Sciences, Stony Brook, NY 11794, USA; Department of Biological Sciences, Hunter College of The City University of New York, USA; Institute for Computational Biomedicine, Weill Cornell Medical College, New York, NY 10021, USA; Wellesley College, Wellesley, MA 02481, USA; Critical Path Institute, Tucson, AZ, USA; DDL Diagnostic Laboratory, 2288 ER, Rijswijk, Netherlands; NSilico Life Science, Nova Center, Belfield Innovation Park, University College Dublin, Dublin 4, Ireland; OTSUKA Pharmaceutical Development & Commercialization, Inc, Princeton, NJ, USA; New York Genome Center, New York, NY 10013, USA

**Keywords:** BioCompute, BioCompute Objects (BCO), high-throughput sequencing (HTS), next generation sequencing (NGS), regulatory review, CWL, FHIR, GAG4H, HL7, Research Objects (RO), Provenance, FAIR data guidelines.

## Abstract

A personalized approach based on a patient’s or pathogen’s unique genomic sequence is the foundation of precision medicine. Genomic findings must be robust and reproducible, and experimental data capture should adhere to FAIR guiding principles. Moreover, effective precision medicine requires standardized reporting that extends beyond wet lab procedures to computational methods. The BioCompute framework (https://osf.io/zm97b/) enables standardized reporting of genomic sequence data provenance, including provenance domain, usability domain, execution domain, verification kit, and error domain. This framework facilitates communication and promotes interoperability. Bioinformatics computation instances that employ the BioCompute framework are easily relayed, repeated if needed and compared by scientists, regulators, test developers, and clinicians. Easing the burden of performing the aforementioned tasks greatly extends the range of practical application. Large clinical trials, precision medicine, and regulatory submissions require a set of agreed upon standards that ensures efficient communication and documentation of genomic analyses. The BioCompute paradigm and the resulting BioCompute Objects (BCO) offer that standard, and are freely accessible as a GitHub organization (https://github.com/biocompute-objects) following the “Open-Stand.org principles for collaborative open standards development”. By communication of high-throughput sequencing studies using a BCO, regulatory agencies (e.g., FDA), diagnostic test developers, researchers, and clinicians can expand collaboration to drive innovation in precision medicine, potentially decreasing the time and cost associated with next generation sequencing workflow exchange, reporting, and regulatory reviews.

## Introduction

Precision medicine requires the seamless production and consumption of genomic information. The National Center for Biotechnology Information’s (NCBI) Database of Genotypes and Phenotypes (dbGaP)[1] and ClinVar[2] illustrate the benefits of genomic data sharing structures such as genome-wide association studies (GWAS). LD Hub, a centralized database of GWAS results for diseases/traits[3], is another example of success. Although the importance of data sharing is established, recording, reporting, and sharing of analysis protocols are often overlooked. Standardized genomic data generation empowers clinicians, researchers, and regulatory agencies to evaluate the reliability of biomarkers generated from complex analyses. Trustworthy results are increasingly critical as genomics play a larger role in clinical practice. In addition, fragmented approaches to reporting impede the advancement of genomic data analysis techniques.

The price of high-throughput sequencing (HTS) decreased from $20 per base in 1990 to less than $.01 per base in 2011[4]. Lower costs and greater accessibility resulted in a proliferation of data and corresponding analyses that in turn advanced the field of bioinformatics. Novel drug development and precision medicine research stand to benefit from innovative, reliable, and accurate -*omics-*based (i.e., genomics, transcriptomics, proteomics) investigation[5]. However, the availability of HTS has outpaced existing practices for reporting on the protocols used in data analysis.

Fast Healthcare Interoperability Resources[6,7] (FHIR) and the Global Alliance for Genomics and Health[8] (GA4GH) capture and communicate genomic information within specific community domains. The Common Workflow Language[9] (CWL) and Research Objects[10] (RO) capture repeatable and reproducible workflows in a domain agnostic manner. The BioCompute framework (https://osf.io/zm97b/) combines these standards via a BioCompute Object (BCO)[11] to report the provenance of genomic sequencing data in the context of regulatory review and research. A BCO is designed to satisfy FAIR Data Principles (findable, accessible, interoperable and reusable)[12], ensuring that data and pipelines are available for evaluation, validation, and verification[11,13,14]. The BCO also meets the NIH strategic plan for data science[15], which states that the quality of clinical data should be maintained at all stages of the research cycle; from generation through the entire analysis process. These characteristics ensure that the BioCompute framework is applicable in any context where scientists are required to report on data provenance, including large clinical trials or the development of a knowledgebase. In the following text, we describe how the BioCompute framework (See Fig. 1) leverages and harmonizes FHIR, GA4GH, CWL, and RO to create a unified standard for the collection and reporting of genomic data.

**Figure 1.**
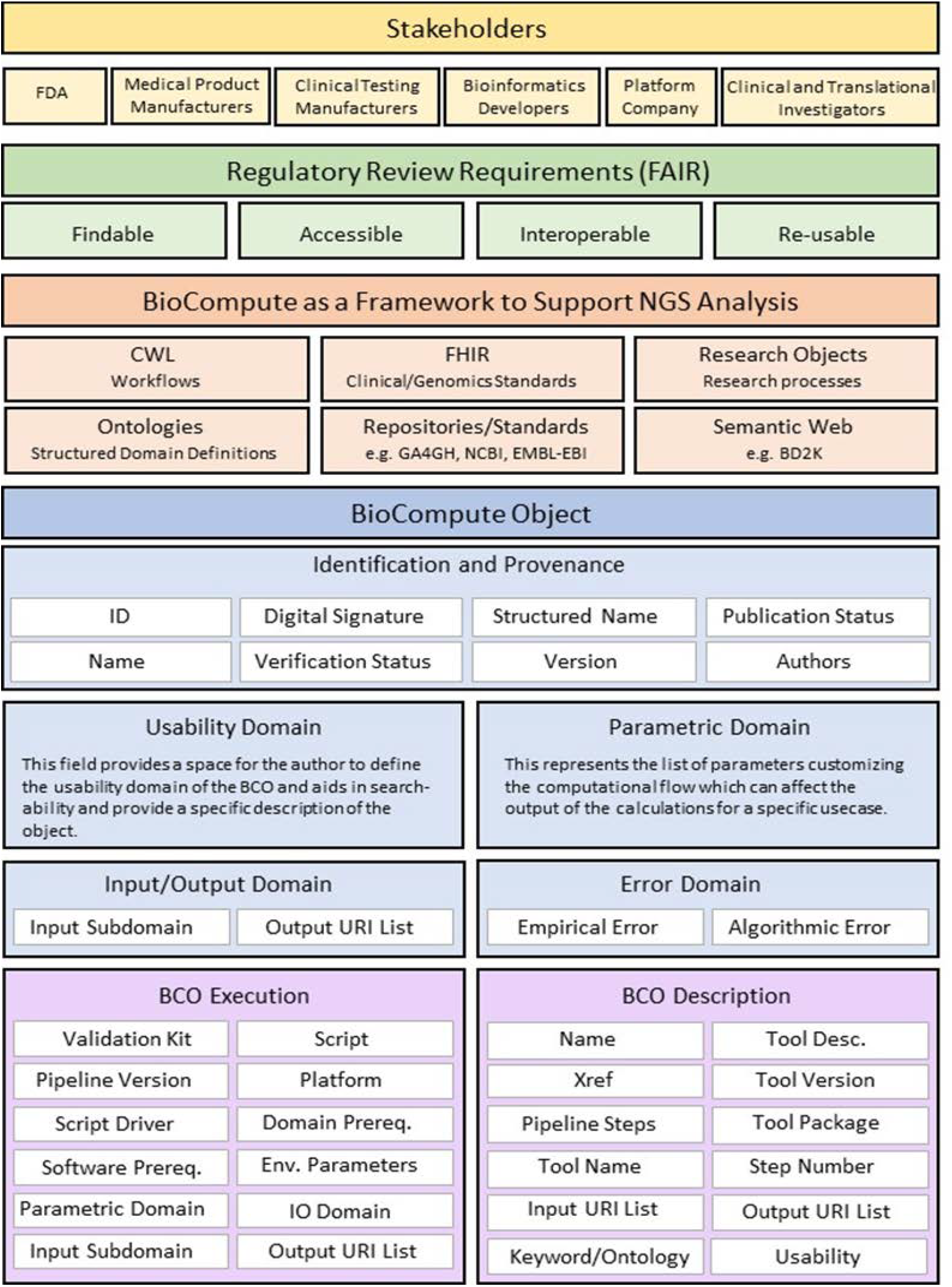
Schematic of BioCompute Object as a framework for advancing regulatory science by incorporating existing standards and introducing additional concepts that include digital signature, usability domain, validation kit, and error domain.

## Background

At a recent Academy of Medical Sciences symposium on preclinical biomedical research, participants identified several measures for improving reproducibility. These include greater openness and transparency, defined reporting guidelines, and better utilization of standards and quality control measures[16]. Compromised reproducibility dissipates resources and hinders progress in the life sciences, as highlighted by several other publications[17,18].

Researchers utilize two types of reproducibility: method reproducibility and result reproducibility[19]. The first type, also called repeatability, is defined by providing detailed experimental methods so others can repeat the procedure exactly. The second type, also called replicability, is defined by achieving the same results as the original experiment by closely adhering to the methods. Method reproducibility depends on a comprehensive description of research procedures. CWL achieves this aim for computational methods by encouraging scientists to adhere to a common language. This facilitates better methods for identification of errors and locating deviances. Research Objects for workflows is built on this concept by using metadata manifests that describe the experimental context, including packaging of the method, provenance logs, and the associated codes and data[20].

Universally reproducible data is an aspirational outcome, but challenges remain. Without widely adopted repeatability and analytical standards for NGS/HTS studies, regulatory agencies, researchers, and industry cannot effectively collaborate to validate results and drive the emergence of new fields[21]. Irreproducible results can cause major delays and be a substantial expenditure for applicants submitting work for regulatory review, and may even be viewed as not trustworthy by third-party verification groups or regulatory agencies. A large number of workflow management systems and bioinformatics platforms have been developed to overcome this barrier, each with their own unique method to record computational workflows, pipelines, versions, and parameters, but these efforts remain disorganized and require harmonization to be fully effective. BioCompute enables clear communication of “what was done” by tracking provenance and documenting processes in a standard format irrespective of the platform or programming language or even tool used.

## Provenance of Data

In order to be reproducible, the origin and history of research data must be maintained. Provenance refers to a datum’s history starting from the original source, namely, its *lineage*. A *lineage graph* can show the source of a datum in a database, data movement between databases or computational processes, or data generated from a computational process. Complementary to data lineage is a *process audit.* This is a historical trail that provides snapshots of intermediary states, values for configurations and parameters, and traceability of stepwise analytical processing[22]. Such audit trails enable an independent reviewer to effectively evaluate a computational investigation. Both types of records gather critical provenance information to ensure accuracy and validity of experimental results. Modern computational workflows produce large amounts of fine-grained but useless trace records, while modern web developments facilitate easy data transformation and copying. Combing and organizing the large volume of resulting material is a daunting challenge. In the molecular biology field alone, there are hundreds of public databases. Only a handful of these retain the “source” data; the remainder consist of secondary views of the source data or views of other publications’ views[23]. Databases are challenged to accurately collect lineage and process records, while also maintaining granularity and ‘black-box’ steps[24,25].

Provenance tracing issues have far-reaching effects on scientific work. Advancement depends on confidence in each of the following: accuracy and validity of the data, process used, and knowledge generated by research. Establishing trust is especially difficult when reporting a complex, multistep process involving aggregation, modeling, and analysis[26]. Computational investigations require collaboration with adjacent and disparate fields to effectively analyze a large volume of information. Effective collaboration requires a solution beyond Open Data to establish Open Science. Provenance must be preserved and reported to promote transparency and reproducibility in complex analyses[27]. Standards must be established to reliably communicate genomic data between databases and individual scientists.

An active community has engaged in provenance standardization to achieve these aims[28], culminating in the W3C provenance specification (PROV)[29]. PROV-O is used by FHIR and ROs and is based on the concept of generating an entity target via an agent’s activity (See Fig. 2). Workflow management systems capture analytic processes while bioinformatic platforms capture analysis details. In combination, they provide a record of data provenance. Few, if any of systems and platforms offer a consistent method to accurately capture all facets of the various roles assumed by an agent who manipulates digital artifacts. BCO encourages adoption of standards such as PROV-O (https://www.w3.org/TR/prov-o/) and ORCID (https://orcid.org/) by defining how to recreate a complete history of what was computed, how it was computed, and by *whom* it was computed and *why* it was computed. Also known as the *provenance domain*, this section of a BioCompute Objects incorporate the Provenance, Authoring and Versioning ontology (PAV, namespace http://purl.org/pav/) to capture “just enough” information to track how data is authored, curated, retrieved, and processed among many specific designations of an “agent”.

**Figure 2.**
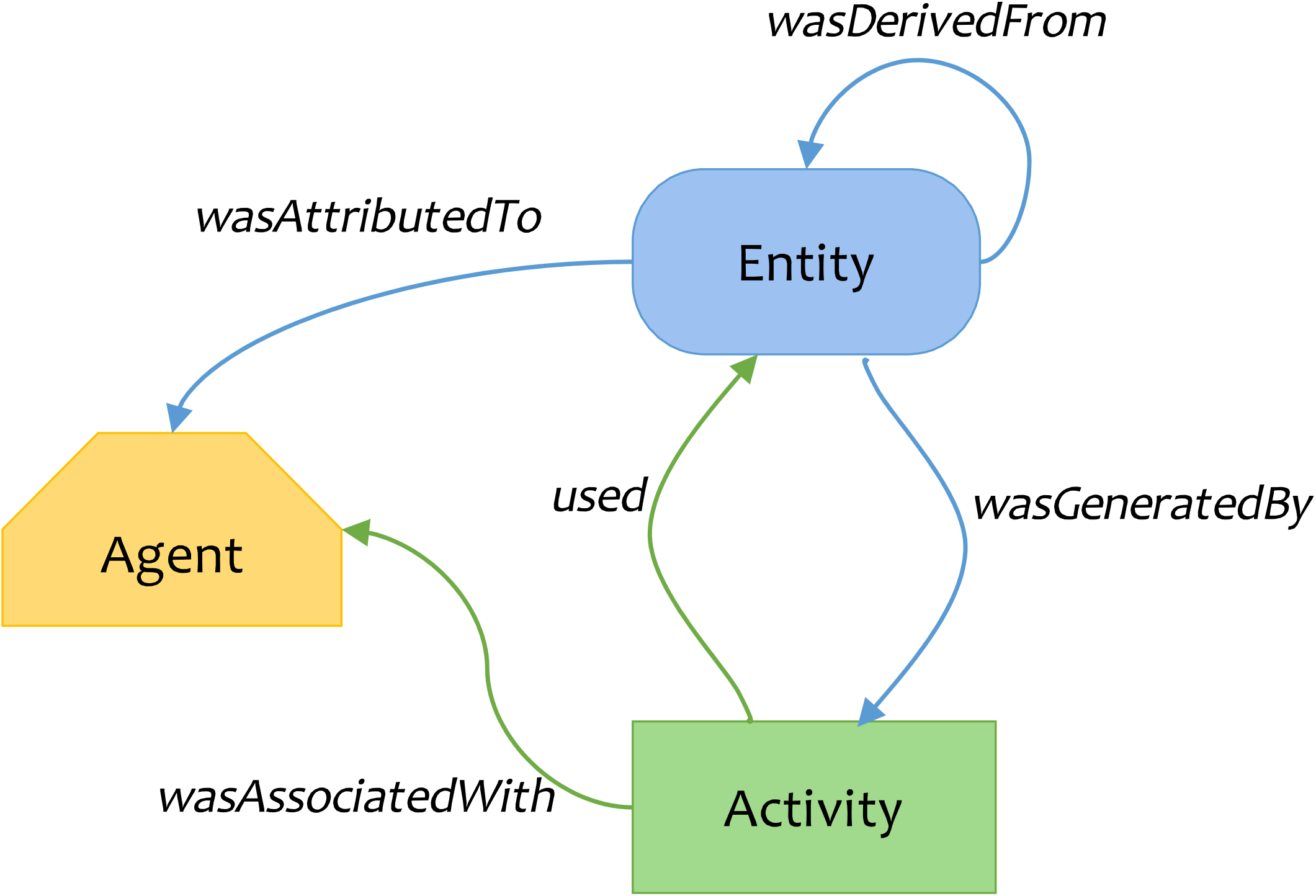
W3C PROV Data Model overview, used in FHIR and RO. Adapted from http://www.w3.org/TR/prov-primer/

## Key Considerations for Communication of Provenance, Analysis, and Results

### Workflow Management Systems

Scientific workflows have emerged as a model for representing and managing complex scientific computations[26]. Each step in a workflow specifies a process or computation to be executed, linked with other steps by the data flow and dependencies. In addition, workflows describe the mechanisms to carry out the steps in a distributed computing environment[26,30]. Documentation of both analytic processes and provenance information is essential for useful reporting[31].

Workflow management systems coordinate the sequential components in an analysis pipeline[30]. They also enable researchers to generate pipelines that can be executed locally on institutional servers, and remotely on the cloud[32]. Cloud infrastructure, high-performance computing (HPC) systems, and Big Data cluster-computation frameworks enhance data reproducibility and portability (see Supplementary File 1). Workflow-centric research objects with executable components are an extension of these management systems[5,20]. Extensive reviews of workflow systems currently in use for bioinformatics have already been published[32-35] and we are not recommending any one over the others. Currently workflow management systems capture provenance information, but rarely in the PROV standard (https://www.w3.org/TR/prov-overview/). Therefore, BCOs rely on existing regulatory standards, that are them-selves are based on PROV standards, like CWL to manage pipeline details, and ROs and FHIR to unify and enhance interoperability.

### Bioinformatics platforms

HTS technology is increasingly relevant in the clinical setting, with a growing need to store, access, and compute more sequencing reads and other biomedical data[36]. The increase in computational requirements have directed the scientists in this space to a call for standard usage methods on integrated computing infrastructure, including storage and computational nodes. This kind of standard will minimize transfer costs and remove the bottlenecks found in both downstream analyses and community communication of computational analysis results[37]. For bioinformatics platforms, communication requirements include (a) recording all analysis details such as parameters and input datasets and (b) sharing analysis details so that others can understand and reproduce analyses.

High-throughput computing (HTC) environments deliver large amounts of processing capacity over long periods of time. These are ideal environments for long-term computation projects, such as those performed for genomic research[38]. Most HTC platforms utilize distributed cloud-computing environments to support extra-large dataset storage and computation, while hosting tools and workflows for many biological analyses. Cloud-based infrastructures also reduce the “data silo” phenomenon by converting data into reproducible formats that facilitate communication (see Supplementary File 1). Additionally, the National Cancer Institute has initiated the Cloud Pilots project, in order to test a distributed computing approach for the multi-level, large-scale datasets available on The Cancer Genome Atlas (TCGA)[39]. Several of the high-throughput[26], cloud-based platforms that have been developed, including HIVE (High-performance Integrated Virtual Environment)[37,40] and Galaxy[41], along with commercial platforms from companies like DNAnexus (dnanexus.com), and Seven Bridges Genomics (sevenbridges.com), have participated in the development of BioCompute. This participation ensures that users while using these bioinformatics platforms would not need to keep track of all of the information needed to create a BCO. Such information will be automatically or semi-automatically collected during the creation and running of a workflow.

The genomic community has come to acknowledge the necessity of data sharing and communication to facilitate reproducibility and standardization[42,43]. Data sharing is crucial in everything from long-term clinical treatments to public health emergency response[44]. Extending bioinformatics platforms to include data provenance, standard workflow computation, and encoding results with available standards through BCO implementation will greatly support the exchange of genomic data analysis methods for regulatory review.

## Regulatory Supporting Standards

Assessment of data submitted in a regulatory application requires clear communication of data provenance, computational workflows, and traceability. A regulatory reviewer must be able to verify that sequencing was done appropriately, pipelines and parameters were applied correctly, to be able to assess that the final result, and critically evaluate the validity of results such as allelic difference or variant call. Because of these requirements, review of any clinical trial or any submission supported with HTS results requires considerable time and expertise. Inclusion of a BCO with a regulatory submission would help to ensure that data provenance is unambiguous and that the bioinformatics workflow is fully documented[11,15,23,45,46].

To truly understand and compare computational tests, a standard method (like BCO) requires tools to capture progress and to communicate the workflow and input/output data. As the regulatory field progresses, methods have been developed and are being continually refined to capture workflows and exchange data electronically[26]. See Figure 3 for BioCompute extensions to HTS analysis that support data provenance and reproducibility.

**Figure 3.**
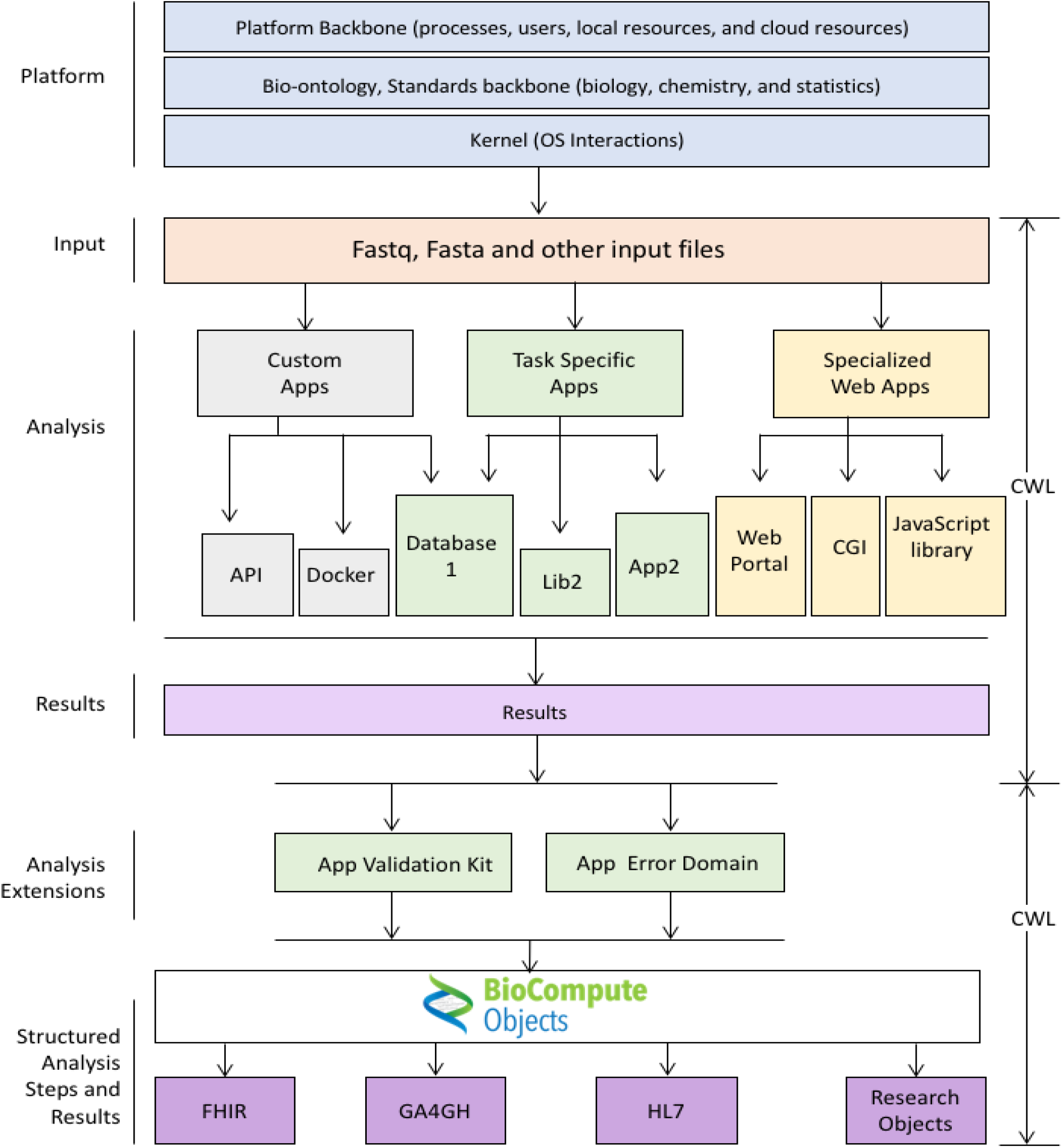
Generic HTS platform schematic with proposed BioCompute Object integrations and extensions.

### Biocompute Objects (BCOs) and Their Harmonizing Efforts

The BioCompute paradigm and Biocompute Objects (BCOs) were conceptualized to capture the specific details of HTS computational analyses. The primary objectives of a BCO is to (a) harmonize and communicate HTS data and computational results as well as (b) encourage interoperability and the reproducibility of bioinformatics protocols. Harmonizing HTS computational analyses is especially applicable to clarify the results from clinical trials and other genomic related data for regulatory submissions.

The BioCompute paradigm is novel in its combination of existing standards with the methodologies and tools to evaluate an experiment both programmatically and empirically. The BCO takes a snapshot of an experiment’s entire computational procedure adhering to FAIR data guidelines. A BioCompute Object is *findable* and publicly *accessible* through the BCO portal, *interoperable* by maintaining the computational context and data provenance which also makes the computational experiment *reusable*. Using this snapshot of the experiment, which can include the range of acceptable experimental results in the verification kit, allows any other user to run the exact experiment and produce the same results. Additionally, through the use of provenance and usability domains, a knowledgeable reviewer can quickly decide if the underlying scientific principles merit approval or further review or reanalysis is required.

### BCO Specification

BCO specification document provides details about the BCO structure (https://github.com/biocompute-objects/BCO_Specification/blob/master/BCO_Spec_V1.2.pdf) (Supplementary File 2). BCOs are represented in JSON (JavaScript Object Notation) formatted text. The JSON format was chosen because it is both human and machine readable/writable. Top-level BCO fields include BCO identifier, type, digital signature, and specification version. The rest of the information are organized into several domains. Below is a brief summary of the type of information present in the various domains. For additional details, please see the latest version of the BCO specification document.

Provenance domain includes fields such as structured name, version, inheritance, status, contributors, license, creation and modification dates. Usability domain provides a clear description of the intended use of the BCO. Extension domain allows users to add content related to other standards that enhances interoperability. Description domain provides details such as keywords, external references and human readable descriptions and sequence of pipeline steps. The execution domain is machine-readable information that can be used to run the entire pipeline. Parametric domain provides information on parameters that were changed from default and the input and output domain provides links to input and output files. The error domain includes details that can be used to verify if a particular BCO has been used as intended and any errors is within the acceptable range. Error domain along with verification kit (such as *in silico* generated read files with known mutations) of the BCO allows verification of a workflow in different bioinformatics platforms for example.

### BCO Implementation

The BCO stores information relating to every package and every script -- including nuanced data that is often not reported, like version number -- in a human and machine-readable format. A typical workflow analyzing HTS data may have a dozen iterations as different software, packages, and scripts are used, each with their own parameters. Through trial and error, various analyses and parameters are refined, which may yield new insights. If new insights are discovered, a snapshot of the state that produced the result in question can be stored as a BCO. To further increase the probability of successful replication, the BCO can even be verified before it is sent out for replication. In this step, a user verifies that all input parameters across the entire analysis pipeline will generate the same output (which can be tested on different data sets).

The BCO is therefore far more efficient than any existing means of communicating HTS analysis information. For example; a researcher has discovered a specific variant that corresponds to a population being studied in her own country and she is interested in learning the relevance of this variant in other populations. By using a BCO, her entire analysis can be quickly understood and repeated by her international colleagues, working with their own data sets, with high confidence in the ability to compare all of the results. If the researcher then uses her data for a clinical trial that is subsequently submitted to the FDA for regulatory review, she can be confident that all of the necessary details are included to successfully repeat her analysis. The final result is a transparent, efficient process that may substantially reduce the amount of time, money, and potential confusion involved with clarifying any details that might otherwise have been overlooked.

In order to form a more cooperative community we have migrated all of the BioCompute development to GitHub (https://github.com/biocompute-objects), and are following the “Open-Stand.org principles for collaborative open standards development (https://open-stand.org/about-us/principles/). The BioCompute specification document has been published as a markdown file so that comments/issues can be addressed using the GitHub issue tracking system. The GitHub organization is setup to have a separate repository for each of the use-case examples or implementations. Each of these will be able to link back to the specification, encouraging parallelized development.

### BCO use case: ReSeqTB-UVP

Several BCO examples are available in repositories under the BioCompute organization on GitHub (https://github.com/biocompute-objects/). As an example, the Unified Variant Pipeline BioCompute Object (UVP_BCO) is described. The complex UVP_BCO captures a validated whole genome sequencing (WGS) analysis pipeline, for The Relational Sequencing Tuberculosis (ReSeqTB) Knowledgebase. This knowledgebase is a one-stop resource for curated <u>Mycobacterium tuberculosis</u> (Mtb) genotypic and phenotypic data that have been standardized and aggregated[47]. It is designed as a global resource to rapidly predict Mtb antimicrobial resistance from raw sequence data. The UVP is an Illumina-based consensus next generation sequencing pipeline comprising several bioinformatic tools with defined thresholds designed to annotate and produce a list of mutations (SNPs and indels) in comparison to a reference Mtb genome. The ReSeqTB platform also includes a database containing phenotypic metadata from culture-based testing and molecular probe-based genotypic data, all housed in a cloud environment enabling multi-tiered user access. In addition, a web-based application containing public data is now freely available (www.reseqtb.org).

A publicly available BCO was developed and released for the UVP to standardize and communicate the process for imputing drug resistance profiles using sequence-based technologies. The UVP_BCO for ReSeqTB delineates the key aspects or domains in both human and machine readable JSON file format. The current version of the UVP BCO can be found in the public BCO GitHub (https://github.com/biocompute-objects/UVP-BCO). As multiple bioinformatic pipelines currently exist for Mtb, the BioCompute paradigm will allow users, including regulators, to identify variables and the ways in which they differ between pipelines, while assessing sources of error, in order to standardize and validate HTS results required for patient management.

## Discussion and Conclusions

Robust and reproducible data analysis is key to successful personalized medicine and genomic initiatives. Researchers, clinicians, administrators, and patients are all tied by the information in electronic health records (EHRs) and databases. Current systems rely on data stored with incomplete provenance records and in different computing languages. This has created a cumbersome and inefficient healthcare environment.

The initiatives discussed in this manuscript seek to make data and analyses communicable, repeatable, and reproducible to facilitate collaboration and information sharing from data producers to data users. Without an infrastructure like BCO, increased HTS sequencing creates silos of unusable data, making standardized regulation of reproducibility more difficult. To clear the bottleneck for downstream analysis, the provenance (or origin) of data along with the analysis details (e.g., parameters, workflow versions), must be tracked to ensure accuracy and validity. The development of high-throughput, cloud-based infrastructures (such as DNAnexus, Galaxy, HIVE, and Seven Bridges Genomics) enables users to capture data provenance and store the analyses in infrastructures that allow easy user interaction and creation of BCOs according to BCO specifications described above and in the specification document (Supplementary File 2).

Platform-independent provenance has largely been ignored in HTS. Emerging standards enable both representation of genomic information and linking of provenance information. By harmonizing across these standards, provenance information can be captured across both clinical and research settings extending into the conceptual, experimental methods and the underlying computational workflows. There are several use cases of such work, including submission for FDA diagnostic evaluations, the original use case for the BCO. Such standards also enable robust and reproducible science, and facilitate open science between collaborators, and the development of these standards is meant to satisfy the needs of downstream consumers of genomic information.

The need to reproducibly communicate HTS computational analyses and results has led to collaboration among disparate industry groups. Through outreach activities including conferences and workshops, awareness of the importance of standardization, tracking, and reproducibility methods has improved[48,49]. Standards like FHIR and ROs capture the underlying data provenance to be shared in frameworks like GA4GH, enabling collaboration around reproducible data and analyses. New computing standards like Common Workflow Language (CWL) increase the scalability and reproducibility of data analysis. The BioCompute paradigm acts as a harmonizing umbrella to facilitate human and machine communication, increasing interoperability in fields that use genomic data. Detailed BioCompute specifications (available at: https://github.com/biocompute-objects/BCO_Specification/) can be used to generate BCOs by any bioinformatics platform that pulls underlying data and analysis provenance into its infrastructure. Ongoing BCO pilots are currently working to streamline the flow with the goal of providing users with effortlessly reproducible bioinformatics analyses. As BCOs aim to simplify the review of data that are essential for FDA approval, these pilots mirror clinical trials involving HTS data for FDA submissions.

Subsequently, the incorporation of a “verification test kit (a test for the range of input values that will still produce the same output values) that evaluates the integrity of the BCO will enable pipelines described by a BCO to be implemented on other servers. The verification test kit would consist of simulated or real HTS data sets where the expected results are known. Each unique implementation of the BCO can then be tested using this verification kit to ensure that it reports the expected results. This work will further the development of the error domain, which describes the observed deviations from the expected results.

We can arrive at values in this domain by repeatedly running the pipeline on the verification kit data and testing the analysis methods deployed. In this way BioCompute can fulfill the goal of communicating exactly what was done, and also what it means, scientifically.

Community involvement has grown to more than 300 contributors, participants, and collaborators from public institutions (including NCI, the FDA, and others), Universities (including George Washington University, University of Manchester, Harvard, and others), and several private sector partners. The BioCompute effort has resulted in two publications, three workshops, and FDA submissions. Efforts to integrate the BioCompute standard into existing platforms mentioned above are underway. The standard has also moved to Working Group status within IEEE (http://sites.ieee.org/sagroups-2791/), and is expected to be an ISO-recognized standard. Finally, publicly accessible databases of crowd-sourced BCO’s that allow researchers or clinicians to reproduce workflows in a variety of contexts are also planned.

## Acknowledgements

We would like to recognize all the speakers and participants of the 2017 HTS-CSRS workshop who facilitated the discussion on standardizing HTS computations and analyses. The workshop was co-sponsored by FDA and GW. The comments and input during the workshop were processed and integrated into the BCO specification document and this manuscript. The participants of the 2017 HTS-CSRS workshop are listed here: https://osf.io/h59uh/wiki/2017%20HTS-CSRS%20Workshop/. This work has been funded in part by FDA (HHSF223201510129C) and The McCormick Genomic and Proteomic Center at the George Washington University.

## Disclaimer

The contributions of the authors are an informal communication and represent their own views.

